# Deep haplotype analyses of target-site resistance locus *ACCase* in blackgrass enabled by pool-based amplicon sequencing

**DOI:** 10.1101/2022.06.22.496946

**Authors:** Sonja Kersten, Fernando A. Rabanal, Johannes Herrmann, Martin Hess, Zev N. Kronenberg, Karl Schmid, Detlef Weigel

**Author notes:** for correspondence (F.A.R.), (D.W.).

## Abstract

Rapid adaptation of weeds to herbicide applications in agriculture through resistance development is a widespread phenomenon. In particular, the grass *Alopecurus myosuroides* is an extremely problematic weed in cereal crops with the potential to manifest resistance in the course of only a few generations. Target-site resistances (TSRs), with their strong phenotypic response, play an important role in this rapid adaptive response. Recently, using PacBio’s long-read amplicon sequencing technology in hundreds of individuals, we were able to decipher the genomic context in which TSR mutations occur. However, sequencing individual amplicons is both costly and time consuming, thus impractical to implement for other resistance loci or applications. Alternatively, pool-based approaches overcome these limitations and provide reliable allele frequencies, albeit at the expense of not preserving haplotype information. In this proof-of-concept study, we sequenced with PacBio High Fidelity (HiFi) reads long-range amplicons (13.2 kb) encompassing the entire *ACCase* gene in pools of over hundred individuals, and resolved them into haplotypes using the clustering algorithm PacBio amplicon analysis (*pbaa*), a new application for pools and for plants. From these amplicon pools, we were able to recover most haplotypes from previously sequenced individuals of the same population. In addition, we analyzed new pools from a Germany-wide collection of *A. myosuroides* populations and found that TSR mutations originating from soft sweeps of independent origin were common. Forward-in-time simulations indicate that TSR haplotypes will persist for decades even at relatively low frequencies and without selection, pointing to the importance of accurate measurement of TSR haplotype prevalence for weed management.

## Introduction

Since the introduction of acetyl co-enzyme A carboxylase (*ACCase*) inhibitors in the late 1980s as herbicides in agriculture, numerous plant species have evolved resistance to these chemicals. The two main mechanisms that led to rapid adaptation are target-site resistance (TSR) and non-target-site resistance (NTSR). TSR is characterized by resistance to high levels of the herbicide, and it originates from gene mutations in the target enzyme causing amino acid changes that preclude the herbicide from binding to the enzyme (Devine and Shukla, 2000). NTSR involves processes to degrade or physically remove the active ingredient from its target, such as enhanced metabolization, decreased absorption or translocation, and sequestration (Devine and Shukla, 2000; Heap, 2014b). There are a number of candidate gene families that contribute to NTSR, including cytochromes P450 monooxygenases, glycosyltransferases or glutathione S-transferases (reviewed in (Gaines *et al*., 2020).

Of particular importance for the European continent, field populations of the grassy weed *Alopecurus myosuroides* have the ability to adapt very rapidly to herbicide treatments, resulting in significant yield losses for farmers (Rosenhauer *et al*., 2013; Varah *et al*., 2019). In fact, widespread resistance to *ACCase* inhibitors in *A. myosuroides* has greatly limited the ability of farmers to effectively control this noxious weed (Délye *et al*., 2010; Rosenhauer *et al*., 2013; Heap, 2014a; Heß *et al*., 2022). The *ACCase* gene in *A. myosuroides*, 12.5 kb in length, is the longest of all three genes with known TSR mutations, with all seven single nucleotide polymorphisms (SNP) that lead to resistance located in the penultimate exon (Délye *et al*., 2005; Xu *et al*., 2014). To date, the occurrence and allele frequencies of *ACCase* SNPs conferring resistance have been investigated in several studies (Délye, Straub, Matéjicek, *et al*., 2004; Menchari *et al*., 2006; Délye *et al*., 2010; Rosenhauer *et al*., 2013), but without considering, at a population scale, the genomic context of the complete *ACCase* gene in which these TSR mutations occur. The other known TSR genes in *A. myosuroides* are *ALS* and *psbA*, with lengths of 1.9 and 1.1 kb, respectively (Gronwald, 1997; Tranel and Wright, 2002).

Pool sequencing offers a cost- and time-saving option for the analysis of many individuals by combining barcoded DNA from multiple samples before sequencing (Ferretti, Ramos-Onsins and Pérez-Enciso, 2013). Over the past decade, pool sequencing has been extensively carried out with Illumina short-reads (reviewed in (Schlötterer *et al*., 2014), including pooled amplicon sequencing approaches for resistance diagnosis assays in multiple species (Délye *et al*., 2015, 2020; Schlipalius *et al*., 2019). Although short sequencing reads generated on the Illumina platform continue to be the gold-standard in terms of raw base quality (greater than 99.9%), they have crucial limitations when compared to long-reads. For instance, due to the limited read lengths (from 50 to 300 bases in paired-end mode), to preserve haplotype information, variant calls have to be phased based on known patterns of linkage disequilibrium when analyzing an entire gene. Phasing accuracy, in turn, also depends on the co-occurrence of heterozygous variants within the range of the library insert size. Sanger sequencing offers the possibility to extend this range to ∼1,300 bases, but larger target regions will still consist of multiple amplicons and thus also require phasing (Sanger, Nicklen and Coulson, 1977; Zhou *et al*., 2000). Moreover, cloning is required if one wants to recover the two phases of a heterozygous individual with confidence (Sanger *et al*., 1980; Délye, Straub, Michel, *et al*., 2004). Long-read amplicons offer many advantages to solve the above mentioned phasing limitations with Illumina short reads and Sanger sequencing (Pollard *et al*., 2018). The most widely used third generation long-read sequencing technologies are from Pacific Biosciences (PacBio) and Oxford Nanopore Technologies (ONT). Unfortunately, in their native form, both suffer from a well-known limitation: a per-base accuracy below 90%, which until recently made reliable variant calling or haplotype determination in Nanopore and PacBio long-read technologies difficult (Korlach, Officer and Biosciences, 2013).

With the introduction of the innovative circular consensus sequencing (CCS) strategy to generate High Fidelity (HiFi) reads in the PacBio sequencing technology (Wenger *et al*., 2019), random sequencing errors can now be corrected, fundamentally improving the most problematic part of long-read technologies (Travers *et al*., 2010; Wenger *et al*., 2019). In addition, by accelerating the polymerase speed and increasing the PacBio sequencing output, sequencing of complete genes of 15-20 kb, and even multiplexing of pooled amplicon samples, has become possible (Travers *et al*., 2010; Wenger *et al*., 2019). In combination with the new clustering software PacBio amplicon analysis (*pbaa*) from Pacbio (Kronenberg, Töpfer and Harting, 2021), this offers completely new perspectives for application of amplicon sequencing in diagnostics.

In this study, we describe a high-throughput PacBio amplicon workflow for pooled samples that can be easily adapted to any gene of interest, independently of the organism. We provide a detailed hands-on laboratory protocol to amplify and long-read sequence loci over 10 kb, as well as analysis recommendations for using the software *pbaa* in pools with up to 200 samples. We demonstrate this workflow with TSR mutations in German field populations of *A. myosuroides*. With the exception of a few low-frequency haplotypes, we were able to recover all individual haplotypes in the pools we tested. Furthermore, we found TSRs resulting from soft sweeps in almost all populations. Using SLiM simulations, we demonstrate that these TSR mutations may persist in field populations for decades to centuries, depending on their starting allele frequencies, even when selection is no longer applied. Therefore, it is strongly advised not to base weed management strategies solely on herbicide applications, but also to integrate mechanical weed management and crop rotation, to keep the incidence of weeds in the field continuously low with a combination of chemical and non-chemical measures.

## Results

### Workflow to sequence and analyze long-read amplicon pools

In a recent study, we sequenced PacBio long-read amplicons of the TSR locus *ACCase* in individuals of 47 European *A. myosuroides* populations (Kersten *et al*., 2021). We discovered a recurrent pattern within field populations of different haplotypes with the same TSR mutation resulting from independent mutation events, as opposed to the same TSR mutation being transferred to other haplotypes by recombination. Characterizing TSR diversity of entire haplotypes to this level of resolution was enabled by two main factors: sequencing of single individuals with HiFi reads and the clustering of these reads to reconstruct both haplotypes in each individual with the *pbaa* tool (Kronenberg, Töpfer and Harting, 2021). However, performing independent DNA extractions and generating long-range amplicons with dual barcodes per individual proved to be both time-consuming and costly. To mitigate these limitations in future studies, we evaluated whether haplotype-level resolution can be achieved by sequencing per-field pools of large numbers of individuals. This is of interest for the further characterization of the origin and evolutionary tempo of herbicide resistance.

For benchmarking purposes, we selected nine populations from our previous study (Kersten *et al*., 2021) and compared the *ACCase* haplotypes determined from 22-24 independently-sequenced individuals to *ACCase* haplotypes inferred from pools of 200 individuals. Each population was sown separately in the greenhouse. Then, we used a paper size template to harvest similar amounts of 4-weeks-old leaf tissue from each plant and pooled them per population prior to DNA extraction (Figure 1a). Next, from 50 ng of DNA (on average ∼65 diploid genome copies per individual in the pool; see Methods), we amplified a 13.2 kb long-range PCR fragment that encompasses the entire *ACCase* coding sequence including introns, plus 585 bp upstream and 364 bp downstream sequences. We used direct dual-indexing per pool, which later allowed multiplexing of all pools on a single SMRT cell. We paid special attention to combine similar amounts of PCR amplicons from all pools, by determining amplicon concentrations with a Qubit fluorometer and an additional gel electrophoresis for cross-validation before combining the pools (Figure 1a). A PacBio amplicon library was then created and sequenced on the Sequel II system (Figure 1b).

**Figure 1.**
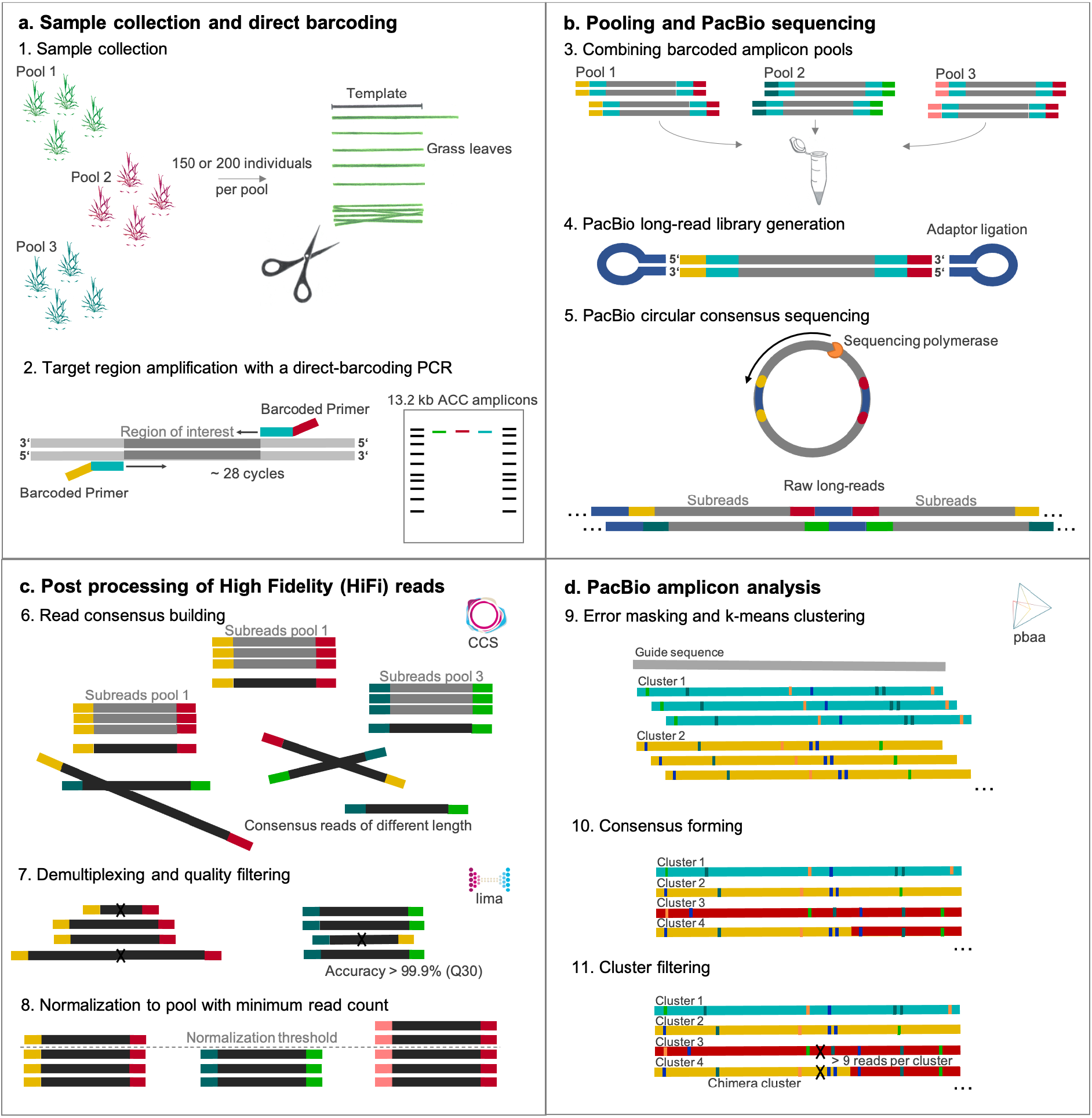
Workflow to generate and analyze long-read amplicons in pools. **a**. Leaf material is collected using a size template to ensure equal sample representation in each pool. Long-range amplicon products are obtained by PCR with direct barcoding to individually tag each pool. Products are visualized by gel electrophoresis for quality control and validation of amplicon concentration measurements. **b**. All population pools are combined in equal amounts in a single tube. A PacBio library is generated and sequenced in circular-consensus mode on a Sequel system. **c**. Computational processing includes read-consensus building, demultiplexing and filtering of raw reads. **d**. *pbaa* clustering is used for variant detection and filtering. The output are *fasta* files listing all haplotypes per population pool and including meta information on read coverage of each haplotype.

The current practice for *de novo* assembly studies, structural variant calling, and amplicon analyses is to start from q20 HiFi reads, that is, reads with an accuracy of at least 99% (Travers *et al*., 2010; Wenger *et al*., 2019). Since we were working with pools consisting of hundreds of individuals, our downstream analysis relied heavily on the precision of each individual read. Therefore, we increased the quality of the input HiFi reads to q30 (≥99.9% accuracy; Figure 1c). To compare haplotype frequencies between populations, we normalized all HiFi reads to the pool with the lowest number of reads, 16,000 reads, corresponding to an average read depth of 40 for each chromosome represented in the sample (200 diploid individuals). Since the most common error types of HiFi reads are indels in homopolymer contexts (Travers *et al*., 2010; Wenger *et al*., 2019), we applied further filters including ‘minimum cluster-read-count 20’ (half the expected depth per single haplotype) and ‘minimum-cluster-frequency 0.00125’, which referred to the fraction of reads to support a true cluster in our datasets (Figure 1d).

### Individuals versus pools - a *pbaa* cluster quality assessment

*pbaa* has been exclusively tested either on single individuals of diploid or polyploid species, or on up to six HLA genes of the same individual (Kronenberg, Töpfer and Harting, 2021). After read-to-read alignment, for each focal read, *pbaa* sorts the alignments in decreasing identity, and retains only the top “n” alignments, which we call a pile. The frequency of each haplotype in the pool affects the parameter choices for *pbaa*. For a perfectly balanced pool, where every haplotype has the same number of reads, the pile size should match the expected haplotype read count. Therefore, to reduce spurious cluster formation, we adjusted the pile size to be about a quarter larger than the expected haplotype read count. The pile is used for error correcting each focal read. If the pile-size is set too high, the pile will contain many cross-haplotype alignments and the haplotype specific variant in the focal read will be corrected away. Similarly, the minimum variant frequency within a pile can affect which variants are masked out. Assuming the pile contains a high fraction of within haplotype alignments, a variant frequency cutoff of 0.4 performs well across a range of parameters. We then compared the resulting haplotypes in pools to haplotypes inferred from individuals of the same populations (Figure 2, Table 1).

**Figure 2.**
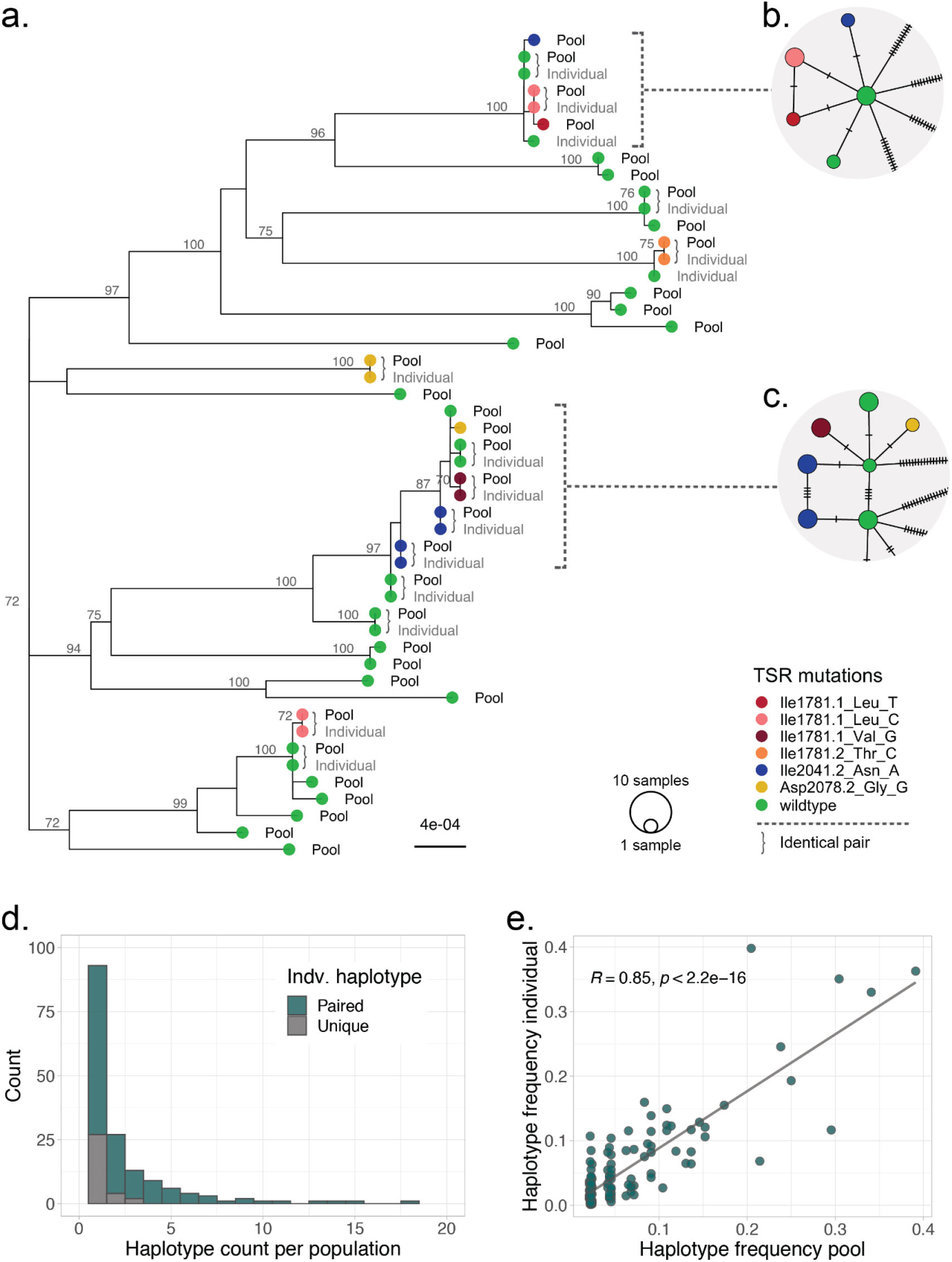
Comparison of haplotypes, as determined with *pbaa*, identified in 24 individuals with haplotypes identified in a pool of 200 *A. myosuroides* samples collected from an agricultural field in Belgian (BE01585). **a**. Maximum-likelihood tree of pool haplotypes and one representative of each cluster inferred in the individual dataset. Tree labels indicate the dataset (Pool vs. Individual). Coloured tree tips show target-site-resistance (TSR) mutations. Curly brackets mark identical haplotype pairs, in which the individual haplotype could be recovered in the pool dataset. **b-c**. Haplotype network representing the corresponding clade in the tree. *pbaa* can successfully recover haplotypes that differ only in one mutation (tick bar). **d**. Haplotype counts per population in the individual dataset. The number of haplotypes that could have successfully identified in the pool dataset is marked in green. Only a fraction of the low abundant ones could not be recovered (gray). **e**. Correlation of haplotype frequencies in the pool dataset versus the individual dataset.

**Table 1.** individual haplotype recovery in pools.

| Population | Number of haplotypes: pool | Number of haplotypes: individuals | Number of correct pairs | Number of correct TSR pairs | Unidentified TSRs in pools | Additional TSRs in pools |
| --- | --- | --- | --- | --- | --- | --- |
| BE01585 | 34 | 15 | 13 (15) | 7 (7) | 0 | 3 |
| DE01467 | 26 | 16 | 14 (16) | 2 (2) | 0 | 2 |
| DE01580 | 31 | 18 | 14 (18) | 3 (3) | 0 | 3 |
| FR01434 | 41 | 21 | 17 (21) | 3 (4) | 1 | 7 |
| FR01729 | 35 | 24 | 19 (24) | 3 (4) | 1 | 6 |
| FR03200 | 24 | 11 | 8 (11) | 3 (3) | 0 | 3 |
| FR07250 | 39 | 18 | 14 (18) | 7 (10) | 3 | 12 |
| NL01505 | 26 | 22 | 19 (22) | 2 (2) | 0 | 0 |
| UK06481 | 34 | 19 | 13 (19) | 4 (5) | 1 | 6 |

The Belgium population BE01585 had a high diversity of TSR haplotypes of independent origin, a classical sign of soft sweeps due to herbicide selection pressure. Among the original 24 diploid individuals, we identified a total of 15 haplotypes, seven of which are haplotypes with TSR mutations (Table 1). In the pool dataset, we successfully recovered 13 of the original 15 haplotypes identified in individuals, including all TSR haplotypes (Figure 2a). The two missing haplotypes in the pool were haplotypes found only in a single individual. Moreover, since the pool contained a larger number of individuals, we could identify 19 additional rare haplotypes, three of which were also TSR haplotypes. Notably, *pbaa* applied to pools was able to correctly resolve haplotypes that differed by a single mutation (Figure 2b-c).

The ability to recover a haplotype in the pools was influenced by its prevalence in the population. All haplotypes that were present in at least four out of 24 individuals were found in the corresponding pool, as were more than 85% of haplotypes present in two or three individuals (Figure 2d). Haplotypes found in only one out of 24 individuals were recovered in 71% of cases. This is most likely a reflection of the experimental design, in which the pools and 24 individuals were drawn from the same seed lots, which contained thousands of seeds, but the 24 individuals were not a subset of the pools of hundreds of individuals. Nevertheless, there was a high correlation (R=0.85, p < 2.2e-16) between haplotype frequency in individuals and in pools (Figure 2e). *pbaa* missed a few TSR haplotypes in the pools compared to the 24 individuals, but in all but one case the analysis recovered additional TSR haplotypes in the pools. As one would expect from the deeper sampling, the number of haplotypes detected in the pools always exceeded the number of haplotypes found among the 24 individuals, from 15% to over twofold (Table 1). Thus, not only did the pools provide valuable, detailed information on the haplotype composition of field populations, but with the identification of up to 12 additional TSR haplotypes, they provided information of importance for resistance monitoring and herbicide use management (Powles and Yu, 2010; Hawkins *et al*., 2018). In addition, the collection of plant pools can constitute a valuable resource for the implementation of standardized epidemiological diagnostic methods, essential for monitoring of future resistances (Comont and Neve, 2021).

### Haplotype clustering reveals evolutionary context

We employed our pool approach to survey TSR haplotype diversity in a German-wide contemporary collection of agricultural fields, for which seeds had been harvested in the year 2019. We selected 64 *A. myosuroides* populations collected in fields of winter annual crops: 49 populations that showed wide-spread resistance to the *ACCase* inhibiting herbicides Axial® (active ingredients 50 g l^-1^ of pinoxaden and 12.5 g l^-1^ cloquintocet-mexyl), 13 populations with an incidence of Axial® resistance below 10%, and two organic fields without a recent history of herbicide application (Data S1). Seventeen farmers were represented with multiple fields. We used the same workflow as described above (Figure 1), but using pools of 150 individuals. To make results comparable across populations, the resulting HiFi reads were normalized to 5,300 reads per pool, corresponding to an average read depth of 17.6 per chromosome (150 diploid individuals). The reads were filtered for ‘minimum cluster-read-count 9’ and ‘minimum-cluster-frequency 0.0017’, which led to an average number of 25 clusters per population (range 15 to 35).

Conventional SNP calling approaches have typically been used for variant calling and analysis of pooled data (Schlötterer *et al*., 2014), but they generally ignore the underlying genomic context. Based solely on allele frequencies, we can estimate the abundance of each TSR mutation (Figure 3a), but we do not know in which context they emerged (Figure 3b). To further assess the accuracy of the pooled clustering approach, we compared the TSR haplotype frequencies with allele frequencies from a conventional SNP calling approach (Figure 3a-b). Pearson correlation coefficients were highly significant and ranged from 0.85 to 1 for all six TSR mutations, with only a few low-frequency *pbaa* clusters not captured, most likely due to the minimum read filter of 9 (Figure 3c).

**Figure 3.**
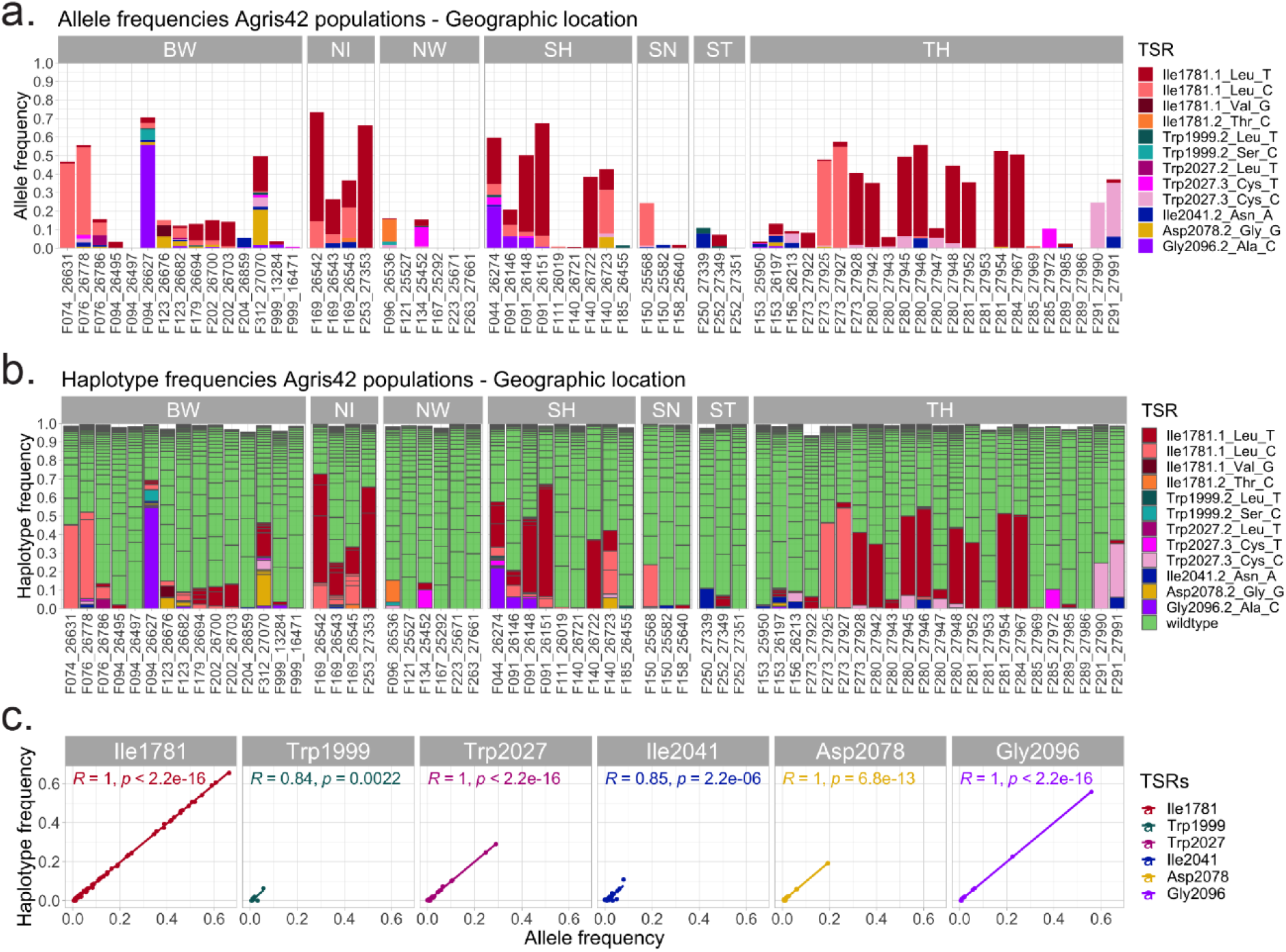
Comparison between conventional single-nucleotide-polymorphism (SNP) mapping and *pbaa* haplotype clustering. **a**. TSR allele frequencies obtained by SNP mapping. Colors indicate different TSR mutations. **b**. Haplotype frequencies inferred using *pbaa*. Colors refer to TSR and wild-type haplotypes. **c**. Correlation between allele frequencies and haplotype frequencies summarized per TSR amino acid position. Correlation coefficients and p-values are shown separately in each TSR panel. BW: Baden-Württemberg, NI: Lower Saxony, NW: North Rhine-Westphalia, SH: Schleswig-Holstein, SN: Saxony, ST: Saxony-Anhalt, TH: Thuringia.

The overall most common TSR mutation is Ile1781Leu, which has been reported in previous studies to increase the fitness of individuals in the absence of herbicide selection and therefore could have been already favorable conditions before herbicide application (Wang *et al*., 2010; Délye *et al*., 2013; Du *et al*., 2019). While the number of populations per state was too small to make definitive statements about regional variation, the state with the smallest fields and farms, Baden-Württemberg, had the most diverse set of TSR mutations and haplotypes. However, states also vary in their history of herbicide use, and thus are not that easily comparable. Very few TSR mutations were observed in field populations of North Rhine-Westphalia, Saxony and Saxony-Anhalt, whereas the field populations in Thuringia seemed to be mainly dominated by single TSR haplotypes.

We refer to a hard sweep when a single haplotype dominates in a population. If, on the other hand, multiple adaptive haplotypes in a population increase in frequency at the same time, this is called a soft sweep (Hermisson and Pennings, 2017). In 38 of 55 German *A. myosuroides* populations containing TSR mutations, we can observe the latter phenomenon, confirming our previous results from European populations where herbicide adaptation occurred predominantly via soft sweeps through TSR mutations of independent origin (Kersten *et al*., 2021). We also find a significant proportion of NTSR for the *ACCase* inhibiting herbicide Axial® in this German dataset, as the biotests reveal significantly more resistance than the TSR frequencies can explain (Figure S1b,d). However, the phenotypic resistance to Focus Ultra correlates significantly with the frequency of TSR mutations Ile1781Leu and Asp2078Gly, as reported before (Powles and Yu, 2010) (Figure S1a,c).

### Organically farmed fields show TSRs of independent origin

Among the nine phenotypically sensitive populations included in the study, there were two organically farmed fields that have not been treated with herbicides going back as far as at least 1980, which predates the introduction of *ACCase* inhibitors to the market. In these fields, we found TSR haplotypes at low frequencies, from 0.3% to 2.0% (Figure 4), in agreement with our previous inferences that standing genetic variation is the most likely evolutionary mechanism behind herbicide selection (Kersten *et al*., 2021). This is considerably higher than in a phenotyping-based study of the grass *Lolium rigidum*, the frequency of resistant individuals to ALS inhibitors in untreated populations ranged from 0.001% to 0.012% (Preston and Powles, 2002). The observation of TSR mutations in organic fields without a history of herbicide use is in agreement with the *ACCase* TSR mutation Ile1781Leu having been detected in one out of 685 (0.146%, or 0.073% at the haplotype level) *A. myosuroides* herbarium specimens collected about a hundred years ago (Délye, Deulvot and Chauvel, 2013). Under herbicide selection, strong resistance can develop within a few generations in such populations. This is due to the fact that mutations present as standing genetic variation have raised to certain frequencies and could already more easily establish in the populations (Hermisson and Pennings, 2005). This is further facilitated by high census population sizes, which can rapidly emerge in years with insufficient weed control and therefore provide a large genetic resource for resistance mutations (Menchari, Délye and Le Corre, 2007).

**Figure 4.**
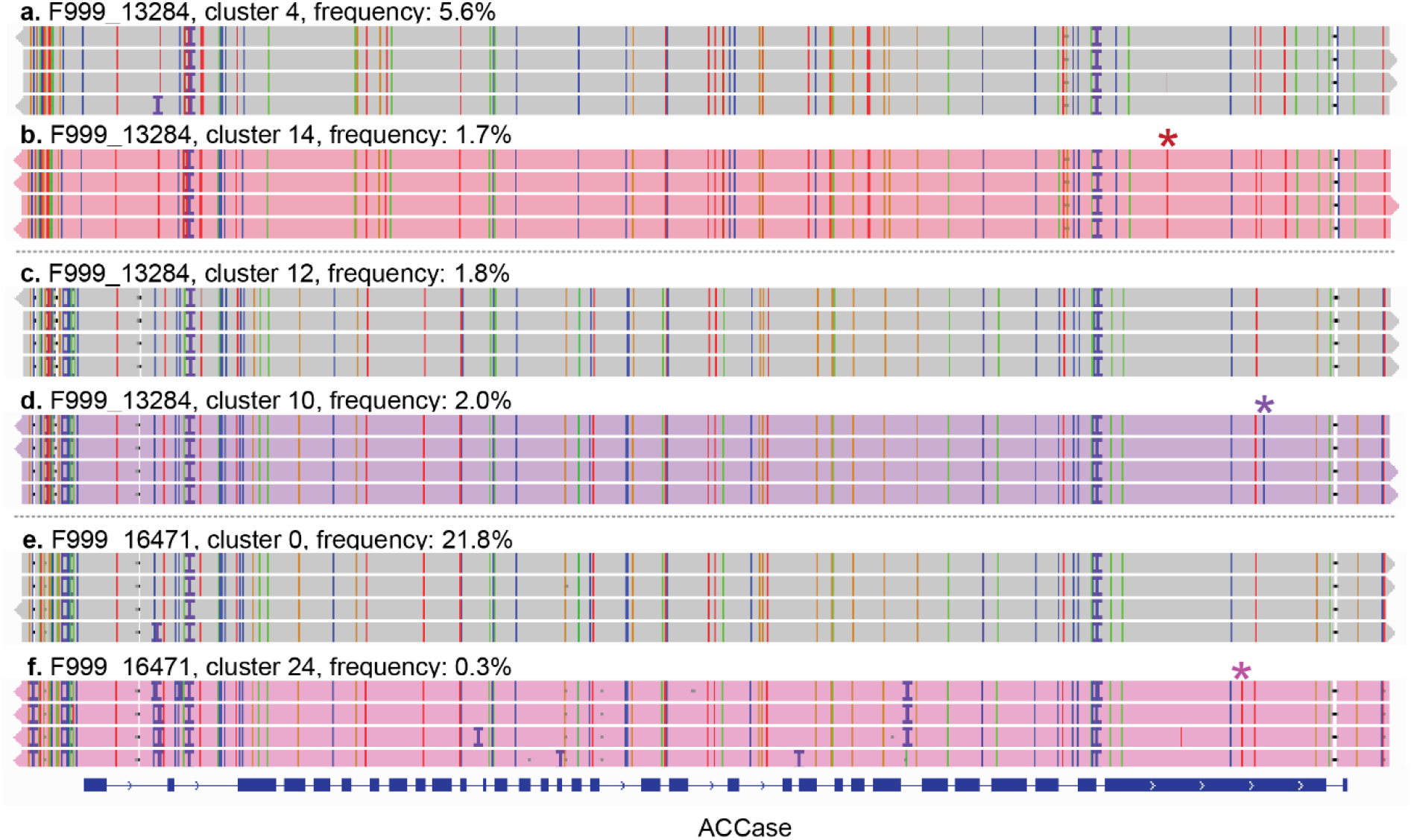
TSR haplotypes and the corresponding wild-type haplotypes from which they arose in organic fields. **a, c, e**. Wild-type haplotypes. **b**. A haplotype with the TSR mutation Ile1781.1Leu_T. **d**. A haplotype with the TSR mutation Gly2096.2Ala_C. **f**. A haplotype with the TSR mutation Trp2027.3Cys_T.

Besides standing genetic variation, another potential source for TSR mutations in these organic fields could be gene flow and seed dispersal by wind, pollen flow, agricultural machinery or wildlife (Colbach and Sache, 2001; Somerville *et al*., 2019). However, in the case of the organically farmed fields, we find not only the TSR haplotypes, but also the corresponding wild-type haplotype, which differs by only one mutation in the entire 13.2 kb amplicon. This is true in all three cases, making it likely that the TSR mutations arose from these wild-type alleles independently in the fields –noticeably, for each field, there are three more pairs of wild-type haplotypes differing by a single mutation from each other. Moreover, we find the corresponding wild-type haplotypes in two out of the three cases in higher frequency than the TSR ones (Figure 4), further suggesting that gene flow as a source is less likely. Instead, their prevalence in the fields serves as a template for different TSR mutations of independent origin.

### TSRs will likely remain in fields for many decades even without selection

Adaptation to new environments is often constrained due to pleiotropic fitness effects in previous conditions (reviewed in (Purrington, 2000). TSR mutations in weed populations represent this special case in agricultural fields, where they become highly beneficial under herbicide application and rise in frequency. However, in the absence of herbicide selection, they have predominantly neutral or even detrimental effects (Menchari *et al*., March, 13 2008; Tardif, Rajcan and Costea, 2006; Vila-Aiub *et al*., 2015; Du *et al*., 2019). At least one *ACCase* mutation, Ile1781Leu, is known to be beneficial under neutral conditions (Wang *et al*., 2010; Délye *et al*., 2013). On the other hand, there have been reports of fitness effects in several TSR mutations in the absence of selection, although quite often the differences do not persist when assessed in realistic field conditions or in competition with other plants (Du *et al*., 2019). Unfortunately, the fitness proxies used, such as biomass, netto assimilation rate, relative growth rate, leaf area ratio, *ACCase* specific activity and plant height, are difficult to compare and to translate into estimates of selection coefficients (Yu *et al*., 2007; Vila-Aiub *et al*., 2015; Sabet Zangeneh *et al*., 2016; Anthimidou *et al*., 2020).

Because herbicide resistance has become such a serious problem in recent decades, it is important to learn whether foregoing herbicide application for certain intervals is sufficient to remove a given TSR mutation from a field population via genetic drift. To tackle this question, we generated forward-in-time simulations with the software SLiM (Haller and Messer, 2019)While most studies focus on few individuals of many populations (Délye, Straub, Michel, *et al*., 2004; Menchari *et al*., 2006), the depth of our pools allows us to assess more realistic haplotype frequencies of TSRs from our empirical dataset (Figure 3). We used high (0.7; Figure 5a,e), intermediate (0.4; Figure 5b,f) and low (0.1 and 0.05; Figure 5c-d,f-g) initial TSR frequencies for our simulations, considering that many TSRs are usually present in the heterozygous state. The simulated selection coefficients ranged from 0 (no detrimental effect in the absence of herbicide selection) to 0.4 (40% fitness cost in the absence of herbicide selection). Within this range, we included the reported selection coefficient estimates for TSR mutations Trp2027Cys and Asp2078Gly, for which under realistic field scenarios seed production was significantly reduced by 20% and 30%, respectively (Du *et al*., 2019). Other parameters, such as effective population size, mutation and recombination rate, were obtained from the literature (Bauer *et al*., 2013; Yang *et al*., 2017; Kersten *et al*., 2021). For each mutation and initial allele frequency, we simulated two different dominance coefficients derived from fitness experiments in *A. myosuroides* (Menchari *et al*., March, 13 2008), an intermediate, codominant coefficient of 0.5 (Figure 5a-d), and a recessive coefficient of 0.25 (Figure 5e-h). We conducted 400 independent SLiM simulation runs per parameter combination and estimated the average number of generations for a TSR to be removed from the population by genetic drift.

**Figure 5.**
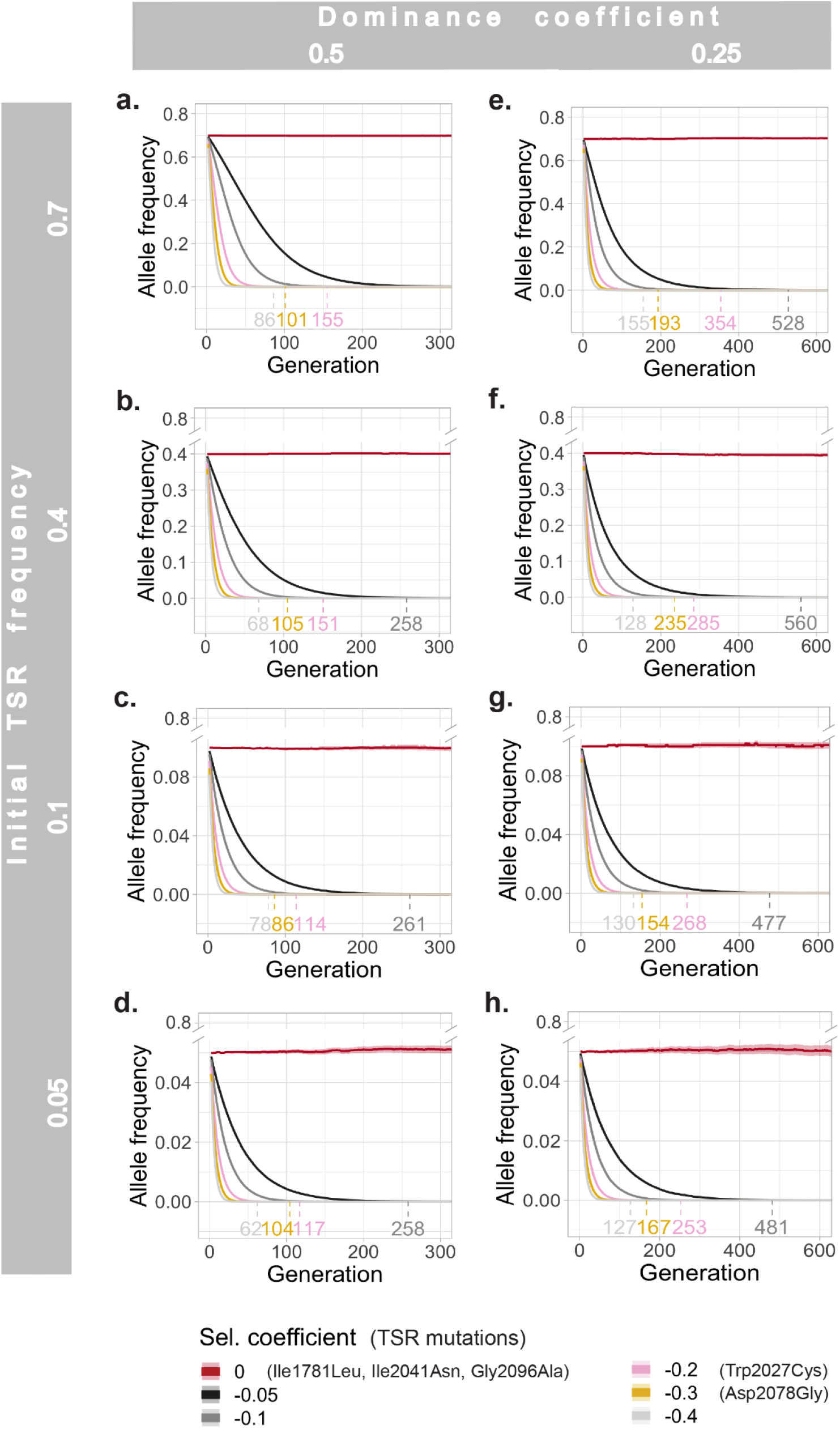
Simulations of the number of generations in which TSR alleles remain in *A. myosuroides* field populations in the absence of selection, assuming different selection coefficients, as estimated from fitness experiments (Menchari *et al*., March, 13 2008; Du *et al*., 2019). While homozygous individuals suffer the full consequences of deleterious TSR mutations, we simulated two different dominance coefficients for heterozygous allele states: an intermediate codominance of 0.5 (**a-d**) and a more recessive coefficient of 0.25 (**d-h**). Means and 0.95 confidence intervals per parameter combination are shown.

The SLiM simulations indicated that under the best-case scenario, with a low initial allele frequency (0.05), a strong deleterious selection coefficient (−0.4), and codominance (0.5), it would take, on average, 62 generations until the TSR mutation is lost (Figure 5d). Unfortunately, farmers often recognize a resistance problem in their fields only once the TSR mutations have already risen to high frequency. Furthermore, most TSR mutations do not seem to reach such a strong deleterious fitness effect (Menchari *et al*., March, 13 2008; Du *et al*., 2019). For example, under a milder selection coefficient of -0.1 (still below what has been reported for Trp2027Cys; Du *et al*., 2019), codominant mutations would persist in a non-treated field for up to 351 generations, and up to 528 generations when more recessive. Since these numbers of generations are mostly beyond the lifespan of a farmer, not to mention the economical loss incurred by a field being fallow for decades, additional measures need to be taken to manage and prevent herbicide resistance.

## Discussion

Pool sequencing of amplicons with PacBio HiFi reads is a cost-effective method for sequencing thousands of samples while preserving haplotype resolution. The *pbaa* clustering software eliminates the need for read alignment against a reference and phasing. Instead, HiFi sequences are clustered directly, preserving the full information contained and reducing reference bias. This opens up new avenues for the discovery of unknown structural variants and genetic diversity. Furthermore, the complete amplicon workflow can be easily established as a high-throughput method for any gene of interest in any organism.

Based on the two population studies we conducted in *A. myosuroides*, Europe-wide and in Germany, we can conclude that herbicide resistance arises independently in different field populations. This puts farmers and consultants in charge to investigate their fields carefully and obtain the status quo of their fields in terms of the resistance situation, because once a resistance mechanism is established in a field population, it is highly unlikely to be lost over the course of a human lifetime, even after herbicide application is stopped. The variation in resistance across the fields sampled in the current study supports the assertion that weed management strategies should focus on the field level, requiring accurate and up-to-date information on the prevalence of herbicide resistance in a given field. A recent survey in Germany found that while only 20% of agricultural fields suffered from high levels of infestation with *A. myosuroides*, resistance to the *ACCase* inhibitor pinoxaden could be detected in 80% of samples (Hess *et al*., 2022). This indicates that successful resistance management requires precautionary control of the census population size of the weed. Management strategies should therefore focus not only on chemical, but also non-chemical measures, such as delayed seeding, moldboard plowing and crop rotation (Moss, Perryman and Tatnell, 2007; Lutman *et al*., 2013).

## Experimental procedures

### European sample collection

The European collection was provided by BASF. Amplicon sequencing data of 22-24 single individuals from 47 populations has been described (Kersten *et al*., 2021). For this study, we selected 9 of those populations containing TSR mutations, re-sowed and sequenced pools of 200 individuals to assess the potential of *pbaa* clustering in pools versus individuals.

### German sample collection and phenotyping

In the course of a Germany-wide herbicide resistance assessment (2019), a collection of *A. myosuroides* seeds on 1,369 agricultural fields was conducted (Hess *et al*., 2022) All samples came from fields sown with winter wheat or triticale in the year of sampling and were screened in a biotest prior to sequencing. Seeds were sown in sandy-loam substrate and treated at BBCH 12/13. Herbicide application was done with 200 liter water in a Research Track Sprayer Generation III using a Teejet-8002-EVS-Nozzle and the corresponding field rate of the herbicides. Two *ACCase*-Inhibitors were used for the screening, Axial® (50 g/l of pinoxaden and 12.5 g/l cloquintocet-mexyl) and Focus® Ultra (100 g/l cycloxydim). A visual assessment of the efficacy was done 21 days after treatment. All plants were screened in two replicates together with well characterized standard populations. 64 samples were later chosen based on the amount of seeds available to conduct further tests, the suitability to form regional clusters, and variations in the degree of efficacy to the tested herbicides. Besides two samples from organic farms all other samples were collected from conventional farms.

### Growth conditions and harvesting

All seeds were sown in standard substrate (Pikiererde Typ CL P, Cat.No EN12580, Einheitserde) and stratified at 4°C in a climate chamber. Then they were transferred to the greenhouse at 22°C with 16 h daylight. For the pilot experiment, we harvested 200 individuals per pool in the European collection. For the German dataset, 150 individuals were collected from each population. To ensure equal representation of all individuals per pool, grass leaves were cut using a 2.5 cm size template (ca. 100 mg leaf material per plant). Care was also taken to ensure that the leaves were of similar width. All pool samples were collected in 50 ml Falcon tubes filled with 4-5 metal beads and ground with a FastPrep tissue disruptor (MP Biomedicals). Then, 1 g of plant leaf powder per pool was transferred to a 2 ml screw cap tube for further processing.

The DNA was extracted with a SPRI-beads based extraction method. The lysis buffer consisted of 100 mM Tris (ph 8.0), 50 mM EDTA (ph 8.0), 500 mM NaCl, 1.3% SDS and 0.01 mg/ml RNase A. For DNA precipitation 5 M potassium acetate was used, followed by two bead-cleanups to purify the DNA (see detailed hands-on extraction protocol: https://github.com/SonjaKersten/TSR_amplicon_pools).

### *ACCase* amplicon generation and PacBio sequencing

To generate the *ACCase* amplicons, we used a direct dual barcoding approach with target-specific primers that had the barcode sequences attached (Data S1). For each PCR, we use 50 ng of DNA as a template. The number of template copies in 50 ng of input DNA of a genome estimated to be 3.6 Gbp (Kersten *et al*., 2021) was estimated to be 13,012 according to the following equation:

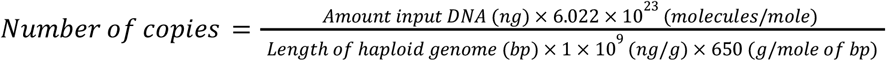

That is, 32.5 and 43.4 copies per haploid genome for pools of 400 and 300 diploid individuals, respectively.

The 13.2 kb long target region was amplified using a master mix reaction with 1 μl Forward indexing primer (5 μM), 1 μl Reverse indexing primer (5 μM), 4 μl 5x Prime STAR buffer, 1.6 μl dNTPs, 0.4 μl Prime STAR polymerase (Takara, R050B), filled up to 20 μl with water in a 2-step PCR reaction with 28 cycles (denaturation: 98°C, 10s; annealing: 68°C, 11 min; final extension: 72°C, 10 min; hold: 4°C). For a quality check, 5 µl of each amplicon pool was visualized on a 0.8% agarose gel and the concentration was determined with a Qubit™ system. Then, all amplicons were pooled equally into a large pool, bead-cleaned and size-selected using a Blue Pippin system (SageScience) with High-Pass Plus 0.75% agarose cassettes, 15 kb (342BPLUS03, Biozym). Only fragments larger than 10 kb were retained. The correct fragment size selection was verified with a Femto Pulse system (Agilent). The PacBio library was created following protocol no. 101-791-800 version 01 (June 2019) with the SMRTbell Express Template Prep Kit 2.0 (part number 100-938-900). Sequel® II loading was performed according to manufacturer specifications with Sequel® II Binding Kit 2.0 and Int Ctrl 1.0 (part number 101-842-900). A detailed hands-on amplicon protocol can be found in Github: https://github.com/SonjaKersten/TSR_amplicon_pools.

### CCS generation and demultiplexing

Pre-processing steps were carried out with PacBio® tools (https://github.com/PacificBiosciences/pbbioconda). This included the generation of circular consensus sequences (ccs) with ccs v6.0.0 with a minimum predicted accuracy of 0.999 (q30), demultiplexing of pools with lima v1.11.0 with parameter settings ‘--ccs –different --peek-guess --guess 80 --split-bam-named --min-ref-span 0.875 --min-scoring-regions 2’, and conversion of the resulting bam to fastq format with bam2fastq v1.3.0.

### *Pbaa*-clustering

Prior to the *pbaa* clustering, we normalized the q30 HiFi reads to the population with the lowest read counts of each dataset with fastqtools v0.8.3 [https://github.com/dcjones/fastq-tools] and indexed each pool with samtools faidx v1.9 (Li *et al*., 2009). In the European collection we used 16,000 reads per pool and in the German collection 5,300 reads. Furthermore, one population in the German dataset did not have enough reads and was, therefore, excluded from further analyses.

The provided guide sequence for the reference-aided clustering approach covered the complete *ACCase* gene sequence and originated from a sensitive plant of a northern Germany reference population (WHBM72 greenhouse standard APR/HA from Sep. 2014) (sequence provided in: https://github.com/SonjaKersten/TSR_amplicon_pools).

In the European dataset, *pbaa* clustering was performed with --min-read-qv 30 --max-alignments-per-read 16000 --max-reads-per-guide 16000 --pile-size 50 --min-var-frequency 0.4 --min-cluster-read-count 20 --min-cluster-frequency 0.00125. In the German dataset, we used the following adjusted parameters: --max-alignments-per-read 5300 --max-reads-per-guide 5300 –pile-size 25 –min-var-frequency 0.4 --min-cluster-read-count 9 --min-cluster-frequency 0.0017 [https://github.com/PacificBiosciences/pbAA]. Finally, to extract the consensus sequences generated in the clustering step, including meta information of each haplotype, and re-orient them –when necessary– in the forward orientation, we used a homemade script, which can be found in the dedicated GitHub for this study (https://github.com/SonjaKersten/TSR_amplicon_pools).

### *pbaa* validation in the European dataset

All clusters inferred by *pbaa* in the pools and all unique haplotypes from the individuals were combined into a joint fasta file per population. MAFFT v7.407 was used for the multiple alignments (--thread 20 --threadtb 10 --threadit 10 --reorder --maxiterate 1000 --retree 1 --genafpair) (Katoh and Standley, 2013) and PGDSpider v2.1.1.5 to transfer the multiple alignment fasta file into a nexus formatted file (Lischer and Excoffier, 2012). The maximum-likelihood (ML)-tree was generated with RAXML-NG v0.9.0 using the GTR+G model and 1,000 bootstraps (Kozlov *et al*., 2019). The minimum spanning network was inferred and visualized with POPART v.1.7 (Leigh and Bryant, 2015). The TSR information for the coloring of the haplotype tree and network was retracted from a classical alignment of the *pbaa* clusters to the *ACCase* reference gene (see section: TSR annotations). The resulting VCF was loaded and manipulated in R to annotate the ML-tree and minimum spanning network. Used R packages can be found in Table S1.

Based on the multiple alignments per population, haplotypes in the pool and individual datasets were counted with the R package ‘haplotypes’ (https://cran.r-project.org/web/packages/haplotypes/haplotypes.pdf) and summarized in Table 1. Haplotype frequencies were calculated with homemade R scripts and the correlations of individual and pool haplotype frequencies were calculated and visualized using the packages ‘ggpubr’ (https://github.com/kassambara/ggpubr/) and ‘ggplot2’ (Wickham, 2016).

### Comparison of conventional SNP mapping with *pbaa* clustering in the German dataset

For the conventional alignment and SNP calling the reads of each pool were aligned to the *ACCase* reference sequence with *pbmm2* [https://github.com/PacificBiosciences/pbmm2]. All resulting bam files were merged, sorted and indexed with samtools v1.9 (Li *et al*., 2009). SNP calling was performed with freebayes v1.3.2 (freebayes -f $REF --min-mapping-quality 20 --min-alternate-fraction 0.005 --pooled-continuous --report-monomorphic) (Garrison and Marth, 2012). All single vcfs were compressed, indexed and merged using tabix v0.2.5 (Li, 2011). To extract the allele depth (AD) and total depth (DP) information, the multiallelic positions were split into multiple rows with bcftools v1.9-15-g7afcbc9 ($bcftools norm -m -any -Oz) (Danecek and McCarthy, 2017), followed by converting the variants into a table using the VariantsToTable function from GATK 4.1.3.0 (Van der Auwera *et al*., 2013). The table was loaded and manipulated in R version 3.6.1 (Team, 2018) and allele frequencies were plotted with the R package ‘ggplot2’ (Wickham, 2016).

### TSR mutation annotation

To annotate the clusters generated with the *pbaa* clustering approach with the TSR information the single cluster fasta files were transferred to *fastq* files with Fasta_to_fastq [https://github.com/ekg/fasta-to-fastq]. Afterwards the *fastq* files containing a single read representing the corresponding cluster were aligned to the *ACCase* reference with minimap2 v2.15-r913-dirty (Li, 2018). The resulting bam file was sorted and indexed with samtools v1.9 (Li *et al*., 2009) and the read groups were adjusted with Picard’s v2.2.1 function AddOrReplaceReadGroups (RGID=$SAMPLE RGLB=ccs RGPL=pacbio RGPU=unit1 RGSM=$SAMPLE) [http://broadinstitute.github.io/picard/]. Variant calling was performed with the HaplotypeCaller from GATK 4.1.3.0 (-R $REF --min-pruning 0 -ERC GVCF), followed by GenotypeGVCFs with standard settings (Van der Auwera *et al*., 2013). Variants in the resulting vcf were annotated with SnpEff v4.3t (Cingolani *et al*., 2012).

### Organic fields with TSR mutations of independent origin

The TSR information of the organic fields was extracted from the previously described haplotype clustering in the German dataset. The clusters in the bam-files were coloured with *pbaa* v.1.0.0 bampaint and visualized in the Integrative Genomics Viewer IGV_2.11.9 (Robinson *et al*., 2011).

### SliM simulations

We performed forward simulations with SLiM v3.4 (Haller and Messer, 2019) under Wright-Fisher model assumptions to determine the number of generations that TSR mutations persist in agricultural fields without being under selection. A population size of 42,000 individuals was assumed, following calculations from the previous publication Kersten *et al*. 2022 (Kersten *et al*., 2021). Similarly we adopted the mutation rate of 3.0 × 10^−8^ (Yang *et al*., 2017) and genome-wide average recombination rate 7.4 × 10^−9^ (Bauer *et al*., 2013) from maize, a diploid grass with a comparable genome size. We modeled a range of selection coefficients (s_i_) from 0 to -0.4, covering values retracted from literature (Du *et al*., 2019). We used two dominance coefficients (h_i_) 0.5 and 0.25 for the TSR mutations as reported in *A. myosuroides (Menchari et al*., *March, 13 2008)*. The fitness model for individuals carrying a homozygous TSR mutation is 1 + s_i_, and for a heterozygous one is 1 + h_i_ * s_i_. Initial haplotype frequencies were extracted from our empirical pool data and set to 0.05, 0.1, 0.4 and 0.7. We performed 400 independent SLiM runs per parameter combination and calculated the mean values and the 0.95 confidence intervals in R with the package ‘rcompanion’ (https://rcompanion.org/handbook/). Visualization was done with ‘ggplot2’ (Wickham, 2016).

## Data availability statement

Raw PacBio subreads will become accessible in the European Nucleotide Archive (ENA; https://www.ebi.ac.uk/ena/browser/home) under project accession number PRJEB53650 upon publication. HiFi q30 reads for each pool can be found at https://keeper.mpdl.mpg.de/d/4ad572a2aab54299a5a2. Experimental protocols, SLiM simulations and custom scripts to reproduce the analyses in this study are deposited in GitHub (https://github.com/SonjaKersten/TSR_amplicon_pools).

## Acknowledgments

We thank Andreas Landes and Jens Lerchl (BASF SE) for providing the Europe-wide populations of *A. myosuroides*, Angela Kuttler and Jakob Keck for help with the sowing and leaf material sampling in the greenhouse, and Christa Lanz (MPI for Biology Tübingen) for assistance with the PacBio amplicon library preparation and the Sequel II loading.

## Author contributions

Conceptualization, F.A.R.; Investigation, S.K. with support from F.A.R., J.H., M.H.; Software, Z.N.K.; Formal Analysis, S.K.; Resources, J.H., M.H.; Writing – Original Draft, S.K.; Writing – Review & Editing Preparation, S.K., F.A.R., D.W.; Visualization, S.K.; Supervision, F.A.R., K.S., D.W.; Funding Acquisition, D.W.

## Supporting information

**Figure S1.**
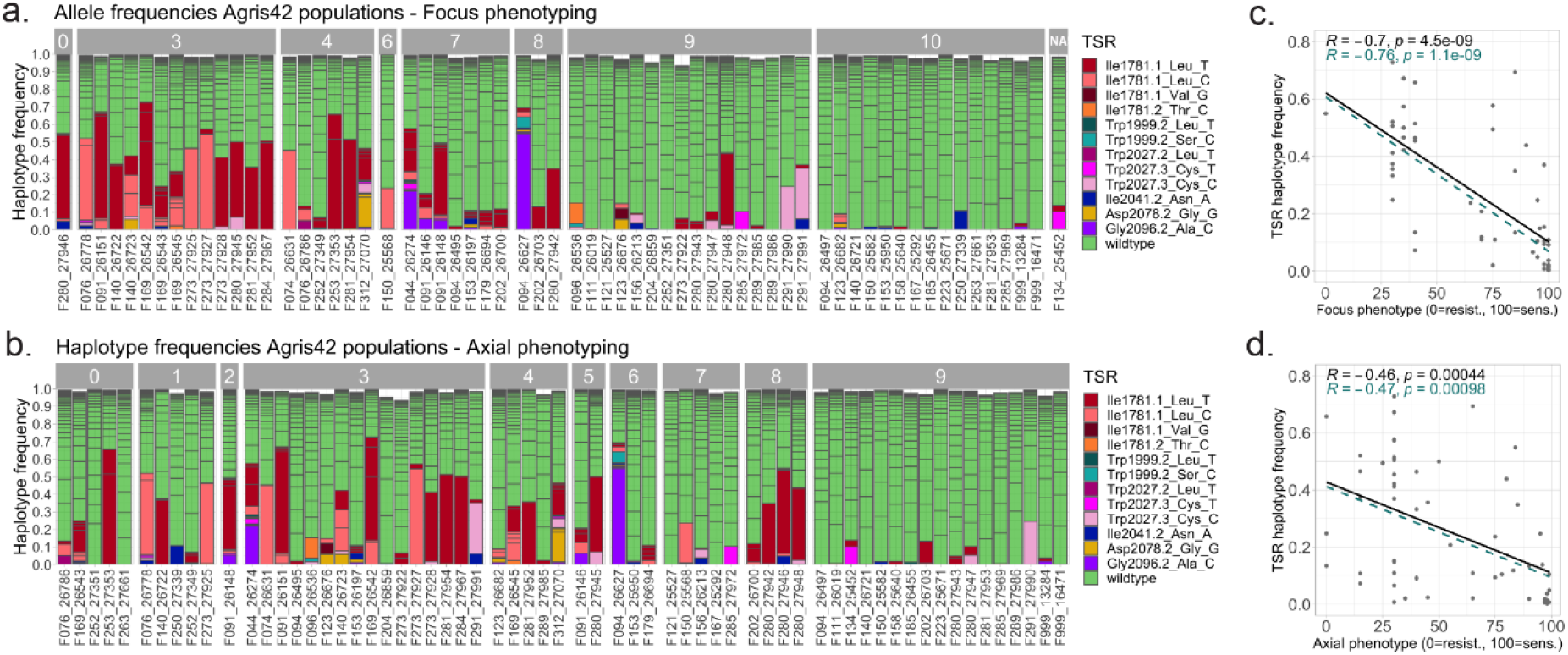
Correlations between TSR haplotype frequencies and phenotyping with ACCase inhibitors. Haplotype frequencies were inferred using *pbaa*. Colors refer to TSR and wild-type haplotypes. **a**. Bins represent the remaining efficiencies of the herbicide Focus Ultra (Bin 0: 0 - 10% efficiency which means 90 - 100% survivor plants, Bin 9: 90 to 99% efficiency, Bin 10: 100% efficiency with 0 % survivor plants). Correlation coefficients and p-values for TSR haplotypes and their respective phenotypes are shown in the panel on the right. **b**. Bins represent remaining efficiencies of the herbicide Axial® (Bin 0: 0 - 10% efficiency, Bin 9: 90 to 99% efficiency, Bin 10: 90 - 100% survivors). **c.-d**. Correlation coefficients and p-values for TSR haplotypes and their respective phenotypes are shown in the corresponding panels on the right. Black line shows the correlation of all TSR mutations, the green line only the TSR mutations with reported resistance to the respective herbicides (summarized in (Powles and Yu, 2010), Table 3).

**Table S1.** R-packages used for data manipulation and visualization.

| **Package name and version** | **Reference** |
| --- | --- |
| dplyr 1.0.2 | (Wickham *et al.*, 2020) |
| ggplot 3.3.2 | (Wickham, 2016) |
| ggpubr 0.4.0 | Kassambara, 2020 (ref<br>(https://github.com/kassambara/ggpubr/) |
| ggtree 1.16.6 | (Yu *et al.*, 2017) |
| haplotypes 1.1.2 | Aktas, 2020 (ref<br>(https://cran.r-project.org/web/packages/haplotypes/haplotypes.pdf) |
| plyr 1.8.6 | (Wickham, 2011) |
| rcompanion 2.4.1 | Mangiafico, 2016<br>(https://rcompanion.org/handbook) |
| reshape 0.8.8 | (Wickham, 2007) |
| tidyr 1.1.2 | Wickham, 2020 (ref<br>(https://github.com/tidyverse/tidyr/) |
| tidyverse 1.3.0 | (Wickham *et al.*, 2019) |
| treeio 1.18.1 | (Wang *et al.*, 2020) |
| vcfR 1.12.0 | (Knaus and Grünwald, 2017) |

**Data S1**. Phenotyping of German populations, target-specific primers with barcode sequences attached, and barcodes to sample correspondence.

